# Speech intelligibility with various head-related transfer functions: A computational modeling approach

**DOI:** 10.1101/2020.06.10.143792

**Authors:** Axel Ahrens, Maria Cuevas-Rodriguez, W. Owen Brimijoin

**Affiliations:** Hearing Systems Section, Department of Health Technology, Technical University of Denmark, 2800 Kgs. Lyngby, Denmark; University of Malaga, 29016, Malaga, Spain; Facebook Reality Labs, Redmond, WA 98052, USA

## Abstract

Speech intelligibility is known to be affected by the relative spatial position between target and interferers. The benefit of a spatial separation is, along with other factors, related to the head-related transfer function (HRTF). The HRTF is individually different and thus, the cues that improve speech intelligibility might also be different. In the current study an auditory model was employed to predict speech intelligibility with a variety of HRTFs. The predicted speech intelligibility was found to vary across HRTFs. Thus, individual listeners might have different access to cues that are important for speech intelligibility.

## 1. Introduction

Speech intelligibility has been shown to be improved in a scenario with spatially separated target and interferer in comparison to a scenario where they are co-located (Bronkhorst, 2000; Hirsh, 1950). This spatial advantage is commonly referred to as spatial release from masking (SRM). Two mechanisms have been shown to be the main contributors to this spatial benefit: a binaural unmasking (BU) component that arises due to different interaural time differences (ITDs) of target and interferer (Durlach, 1963, 1972; Licklider, 1948) and a signal-to-noise ratio (SNR) component (often referred to as better-ear SNR) that arises due to a direction dependent spectral filtering of the source signals from torso, head and pinna. This filter function is also referred to as the head-related transfer function (HRTF) or as head-related impulse response (HRIR) in the time domain. It should be noted that the spectral filtering gives rise to interaural level differences (ILDs), but the HRTF/HRIR also includes ITDs. Previous studies have investigated speech intelligibility while varying ITDs or ILDs and showed that larger differences in ITDs and ILDs lead to better speech intelligibility (Best *et al.*, 2013; Bronkhorst and Plomp, 1988; Pollack and Pickett, 1958). Culling *et al.* (2004a) examined the role of ITD and ILD cues on speech intelligibility separately and in combination, finding better speech intelligibility performances when both cues were present (HRTF without manipulation). However, when interferers were presented in both hemispheres, the advantage caused by the ITD-only HRTFs was bigger than by the ILD-only HRTFs.

HRTFs have been shown to be highly individual. The individual differences have been shown to be due to anthropometric features such as the shape and the size of the head and the pinnae (Algazi *et al.*, 2001; Watanabe *et al.*, 2007). Listening with non-individual HRTFs has been shown to affect perception (see Xu *et al.*, 2007, for a review). For example, sound source localization in the median plane has been shown to be reduced, while localization on the horizontal plane has shown to not be affected by the HRTF (Møller *et al.*, 1996). However, the effect of HRTFs on speech intelligibility remains unclear. Jakien *et al.* (2017) measured speech intelligibility with individual HRTFs when reproducing the stimuli over loudspeakers as well as with artificial head HRTFs when using headphones. They found worse speech intelligibility with the dummy head HRTF than with individual HRTFs. However, they did not include head-movements in the headphone condition but did in the loudspeaker condition, rendering direct comparison problematic. Cubick *et al.* (2018) investigated the effect of the microphone placement in hearing aids on speech intelligibility, which affects the HRTF, and showed a reduced spatial benefit in normal-hearing listeners with hearing aids in comparison to a condition without hearing aids. While these studies do suggest an effect of the HRTF on speech intelligibility, no studies have examined this effect in a direct and controlled way.

In the present study, it was investigated whether HRTFs measured from various individuals have an effect on the predicted speech intelligibility with a computational binaural speech intelligibility model. Thus, it can be investigated if some listeners might have access to more advantageous cues for speech perception based on their HRTFs. Furthermore, using an auditory model allows to draw conclusions about possible underlying mechanisms, while eliminating cognitive factors that might play a role in understanding speech (Heinrich *et al.*, 2015; Ng *et al.*, 2013; Rönnberg *et al.*, 2016). Particularly it was the intent to investigate whether the BU component or the SNR component have more influence on speech intelligibility with various HRTFs.

## 2. Methods

### 2.1 Binaural speech intelligibility model

We used the computational binaural speech intelligibility model from Jelfs *et al.* (2011) as implemented in the Auditory Modeling Toolbox (Soendergaard and Majdak, 2013). The model predicts the target-to-interferer ratio as the sum of a BU component and the SNR component at the ears. The BU component is based on binaural masking level difference (BMLD) predictions (Culling *et al.*, 2005) and is calculated from the ITDs of the target and the interferers and the interaural coherence of the interferer. The SNR component is calculated from the maximum of the long-term SNR of both ears within gammatone filters. The SNR component is also often referred to as better-ear SNR, but can also be nonzero for a symmetric interferer configuration, where spectral filtering leads to SNR advantages over a co-located configuration, that are the same in both ears. The two components are calculated in 43 gammatone filters and integrated over frequency using the speech intelligibility index (SII) weighting (ANSI, 2017). The model takes binaural room impulse responses for the target and the interferers as input signals, but has also been verified with anechoic HRIRs (Jelfs *et al.*, 2011). Using the impulse responses resembles a situation with stationary noise interferers having the same spectrum as the target speech. When multiple interferers are used, the HRIRs were concatenated as described in Jelfs *et al.* (2011).

### 2.2 HRTF databases

Two HRTF databases were used in this study, the LISTEN (Warusfel, 2003) and the CIPIC (Algazi *et al.*, 2001) database. The LISTEN database contains HRTFs measured from 187 directions for 51 subjects. The horizontal plane is fully sampled in 15° steps. Here, the processed version of the HRIRs was taken where the HRIRs were truncated to a length of 512 samples (sampling frequency 44.1 kHz) and diffuse field equalized. The CIPIC database consists of HRIRs from 45 subjects, measured from 1250 directions. The horizontal plane is sampled at 80°, 65°, 55° on both hemispheres as well as in 5° steps between ±45°. Two of the subjects in the CIPIC database are artificial head measurements on a KEMAR (GRAS Sound & Vibration A/S, Holte, Denmark) with a small or a large pinna. The HRIRs are 200 samples long (sampling frequency 44.1 kHz) and were also diffuse-field equalized.

### 2.3 Experimental setup

The target source was always simulated from the front, i.e., 0° azimuth, 0° elevation. Two conditions were investigated, a single and a two interferer condition. In the single interferer condition a large better-ear SNR is expected, while with symmetric interferers this effect is expected to be reduced. The interferers were simulated at various azimuths either on the right hand side in the single interferer condition or symmetrically on both sides in the two interferer condition. Here, we report the predicted SRM, i.e., the difference between the co-located and the separated interferer configurations.

## 3. Results

### 3.1 Single interferer configuration

Figure 1 shows the model predictions with the various HRTFs from the LISTEN (left column) and the CIPIC (right column) database with respect to the azimuth interferer angles (grey lines). The top row shows the predicted SRM, the middle row the predicted SNR component and the bottom row the predicted BU component. The median SRM (black line) across the HRTFs increases with increasing angular separation between the target and the interferer up to about 75° and 65° for the LISTEN and the CIPIC database, respectively. Beyond this angle the median SRM decreases slightly. However, for some HRTFs the predicted SRM continues to increase, for some it remains constant, and for some it decreases. A similar trend can be seen for the predicted SNR advantage. The BU advantage increases up to an angle of 45°.

**Fig. 1:**
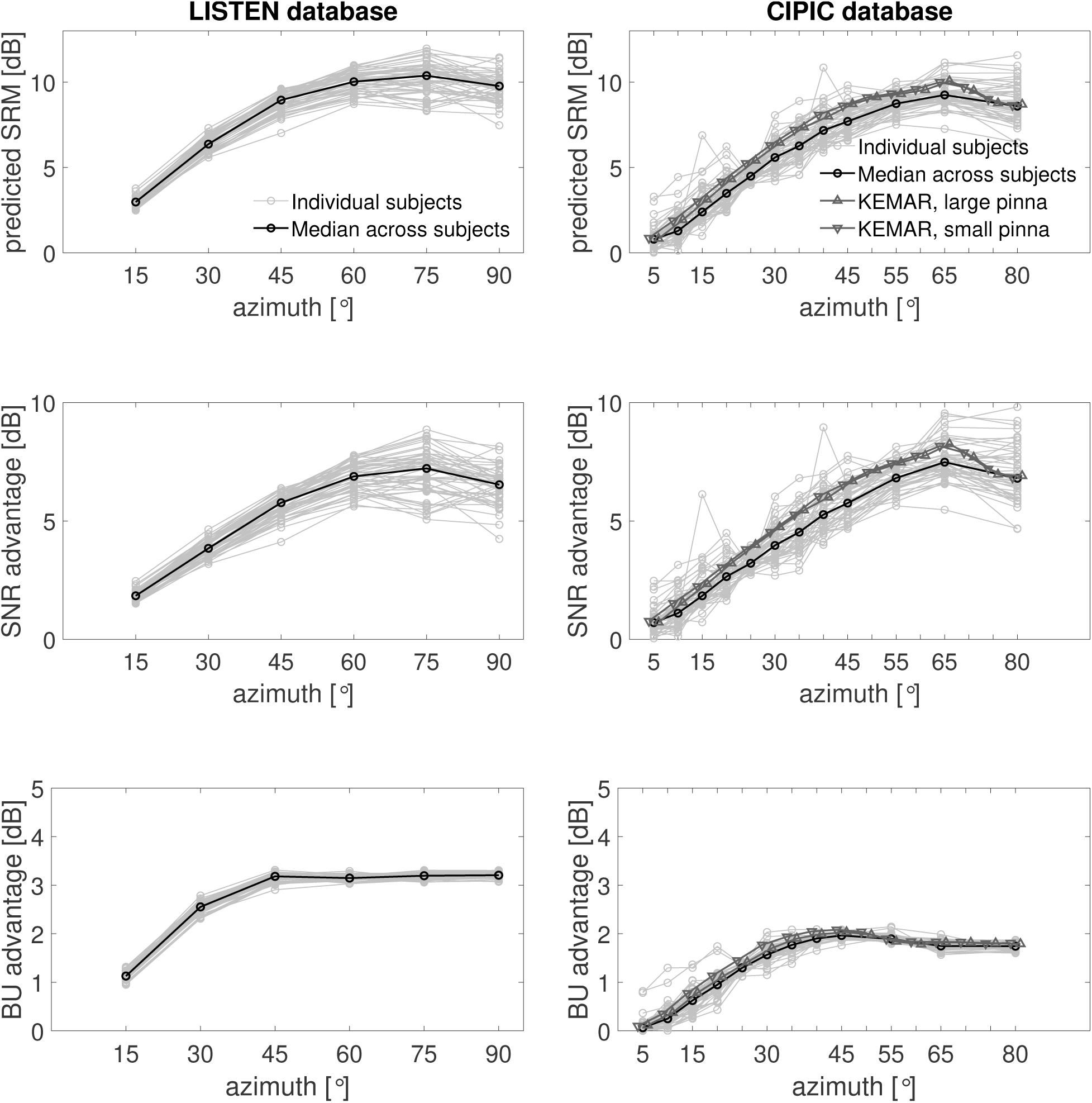
The predicted SRM (top), SNR advantage (middle) and BU advantage (bottom) with respect to the azimuth angle of a single interferer with a target from the front (0°). The grey curves depict individual subjects’ HRTFs from the LISTEN (left) and the CIPIC (right) database. The black curves depict the median calculated over the HRTFs. For the CIPIC database also predictions with artificial head HRTFs are shown (triangles).

The variance of the spatial benefit across HRTFs was found to increase with the separation angle for the LISTEN database but relatively constant for the CIPIC database. The largest difference in predicted SRM was found to be 4 dB for a separation angle of 90° and 5.9 dB for a separation angle of 15°, for the LISTEN and the CIPIC database, respectively. The large difference in the CIPIC database at 15° might be driven by a single outlier. The second largest difference was found at 80° (5.2 dB). To avoid outliers in the data, a percentile analysis has been performed. The range between the 5^*th*^ and the 95^*th*^ percentile was found to be 3.2 dB (at 75°) for the LISTEN and 3.6 dB (at 80°) for the CIPIC database.

To analyse the contribution of the SNR component and the BU component to the variance of the predicted SRM across HRTFs, the R^2^ of a linear regression model was investigated. Therefore, models were fitted either between the SRM and the BU or between the SRM and the SNR. This analysis was performed for the angle with a largest variance only. The correlation analysis between the predicted SRM and the SNR or the BU component of the model revealed that the SNR component alone can account for most variance across HRTFs. The SNR component accounts for 99.7% of the variance in the CIPIC database and 99.5% in the LISTEN database. The BU component accounts for 9.6% in the CIPIC and 1.2% in the LISTEN database.

The predicted SRM was found to be about 2 dB larger for the HRTFs in the LISTEN database than for the ones in the CIPIC database. This difference is mainly driven by the BU component of the model while the SNR component is similar across the two databases.

The CIPIC database also contains two HRTF measurements of an artificial head (KEMAR) with two different pinna sizes. The predicted SRM with the KEMAR HRTFs (dark grey lines) was found larger than the median of thu subjects’ HRTFs. This difference was driven by the SNR component of the model, while the BU component was found similar between subjects and artificial head. The difference between the two pinna sizes was found negligible.

### 3.2 Symmetric interferer configuration

Figure 2 shows the model predictions as in Figure 1 but for symmetric interferers. The median predicted SRM (top panels, black lines) increases with separation angle but less than with a single interferer. The reason is a significantly smaller SNR component (middle panel) due to a reduced better-ear listening advantage. The SNR advantage is close to zero and for some conditions negative. The BU advantage is similar as with a single interferer.

**Fig. 2:**
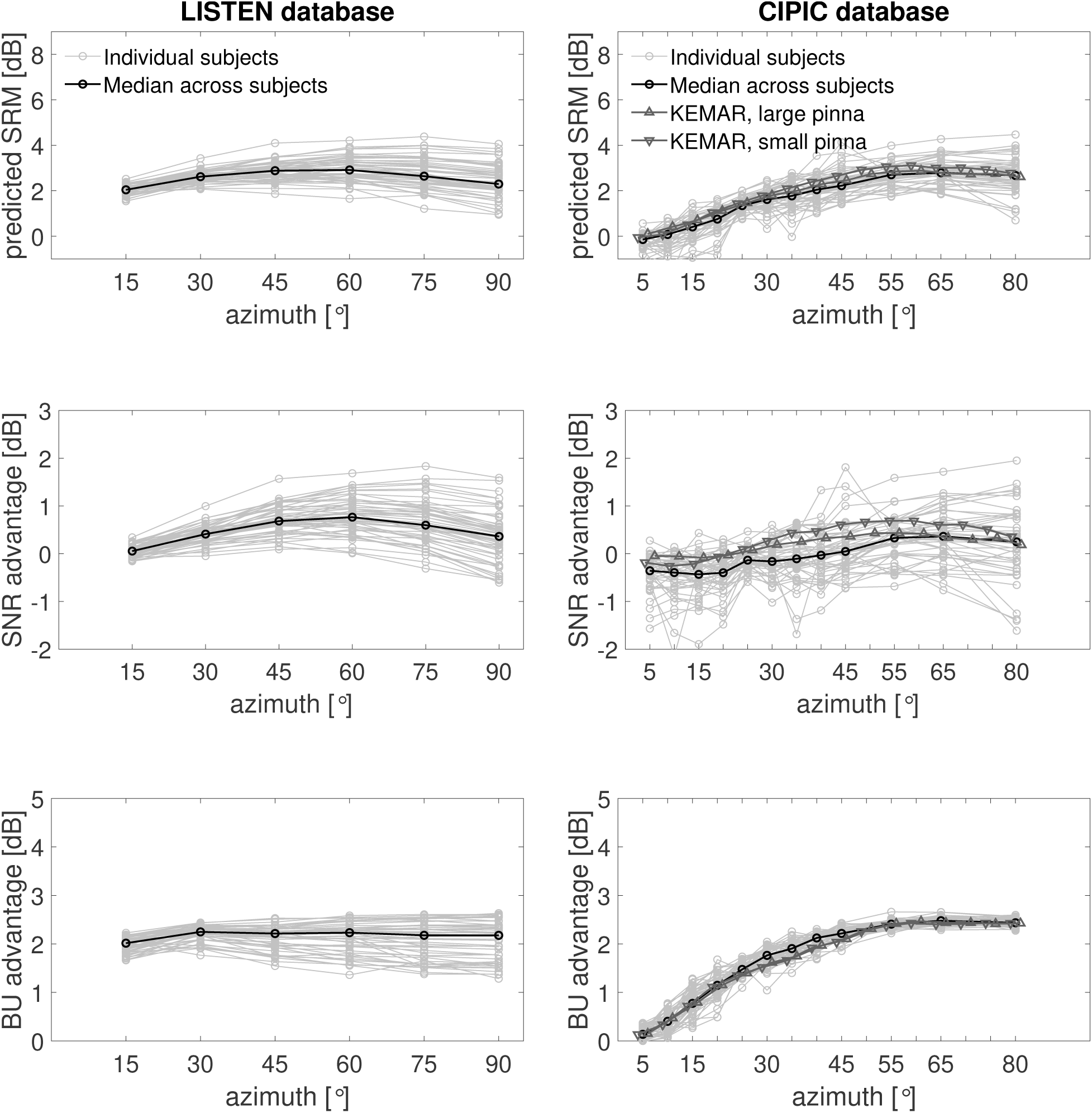
The predicted SRM (top), SNR advantage (middle) and BU advantage (bottom) with respect to the azimuth angle of two symmetric interferers with a target from the front (0°). The grey curves depict individual subjects’ HRTFs from the LISTEN (left) and the CIPIC (right) database. The black curves depict the median calculated over the HRTFs. For the CIPIC database also predictions with artificial head HRTFs are shown (triangles).

The largest inter-HRTF difference was found 3.2 dB (at 75° azimuth) and 3.8 dB (at 80° azimuth) in the LISTEN and CIPIC database, respectively. The range between the 5^*th*^ and the 95^*th*^ percentile was found 2.5 dB at 90° azimuth for the LISTEN and 2.8 dB at 80° azimuth for the CIPIC database.

A correlation analysis using *R*^2^ as in the previous section showed that with symmetric interferers, the SNR advantage alone accounts for 70.4% (at 90° azimuth) and 99.1% (at 80° azimuth) of the variance in the predicted SRM for the LISTEN and the CIPIC database, respectively. The BU component alone can account for 46.4% and 1.2% of the variance, for LISTEN and CIPIC, respectively.

Comparing the two HRTF databases shows a different trend than with a single interferer. For the 15° separation angle between target and interferers, the median predicted SRM is about 1 dB higher with HRTFs from the LISTEN database than from the CIPIC database. The difference is driven by the BU advantage. The SNR advantage in both databases is close to 0 dB. The predicted SRM using the KEMAR HRTFs was comparable to the SRM of the subjects’ HRTFs in the CIPIC database.

## 4. Discussion & Conclusion

In the present study we investigated the influence of various head-related transfer functions (HRTFs) on the spatial release from masking (SRM). We used a computational auditory model and simulated the SRM in various conditions with HRTFs from two publicly available databases. It has been shown that the predicted SRM using a binaural speech intelligibility model is dependent on the HRTF. The variance of the predicted spatial benefit across HRTFs was found larger for wider separation angles between target and interferers than for smaller separation angles. The differences across the HRTFs could mainly be explained by the signal-to-noise ratio (SNR) at the ears while the binaural unmasking (BU) advantage showed less variations. The predicted speech intelligibility with the HRTFs of an artificial head was larger than the subjects’ HRTFs medians in the condition with a single interferer but very similar in the presence of two symmetric interferers. With the artificial head the BU component was closer matched to the subjects’ HRTFs than the SNR component.

The results from the current study are in general agreement with previous speech intelligibility studies. Multiple studies investigated speech intelligibility with a target talker from the front and a single interferer at multiple separation angles on the horizontal plane (Bronkhorst and Plomp, 1988; Peissig and Kollmeier, 1997; Platte and Hövel, 1980; Plomp and Mimpen, 1981). The studies generally found the same trend (see Bronkhorst, 2000, for a summary), in that the largest SRM occurs when the interferer is between 90° to 120°. However, the differences between an interferer at 60° and at 90/120° was small in comparison to the difference between 0° and 60°. These findings are in agreement with the predictions shown in Figure 1. Marrone *et al.* (2008) measured speech intelligibility with a target at 0° and two symmetrically separated interferers and found a plateauing release from masking for separation angles between 45° and 90°, which is in line with the predicted intelligibility results found in the current study (see Figure 2). Similar results were shown by Cosentino *et al.* (2014). The reduction in SRM and particularly in better-ear SNR advantage has previously been shown when interferers are placed in both hemispheres (e.g. Culling *et al.*, 2004b; Hawley *et al.*, 2004).

Here, we entirely relied on predictions from the computational auditory model from Jelfs *et al.* (2011) instead of listener studies. The reason was to only investigate the cues that are available to the auditory system, while avoiding within-listener variations as well as cognitive effects. The risk with a pure modelling approach is that the model does not reflect listener performance. However, Jelfs *et al.* (2011) evaluated the auditory model with various listener studies and showed that the model predictions match the speech intelligibility data well. The most relevant dataset for the current study that they used to evaluate their model was from Bronkhorst and Plomp (1988), where speech intelligibility was measured with a single noise source positioned at various azimuth angles. The results of the model and the measurements both showed an increase of SRM with increasing separation angle. However, the model predicted a lower SRM at 90° than at 60°, while the measured SRM did not show this drop. This drop in SRM at 90° can also be seen in Figure 1 of the current study. Such a drop in intelligibility has been shown previously (Duda and Martens, 1998; Peissig and Kollmeier, 1997). Jelfs *et al.* (2011) argued that the reason for the difference between the model and the data are the use of different HRTFs. This explanation is in line with the findings in the current study. For some HRTFs the 90° drop can be seen, while for others the SRM remains constant or increases.

To our knowledge only one previous study investigated speech intelligibility with human listeners measured with multiple HRTFs (Picinali *et al.*, 2019). They found small but significant effects on speech intelligibility across seven HRTFs from the LISTEN database when using a target talker at 0° and two interfering speech sources at ±90°. Cubick *et al.* (2018) measured speech intelligibility in normal-hearing listeners with and without hearing aids. They showed that hearing aids vary the HRTF as well as reduce the spatial benefit. Employing a similar model as in the current study showed that mainly the better-ear SNR could account for the findings.

The contribution of the SNR component to the variance in predicted speech intelligibility across HRTFs was more pronounced in the condition with a single interferer than with symmetric interferers. The likely reason is that with a single interferer, the ear contralateral to the interferer receives a more favourable SNR than the ipsilateral ear. This effect is commonly referred to as better-ear SNR listening. When the better-ear SNR is large, any variance in HRTFs is likely to contribute relatively more to the SNR component of the model. When the interferers are symmetric, on the other hand, the SNR component is comprised of only differences in the spectra across the ears. We argue that this distinction underlies the observed difference in predicted intelligibility across the symmetric versus asymmetric conditions. However, in both interferer conditions (with symmetric and asymmetric interferers) the variance across HRTFs was mainly driven by the SNR component. Thus, the HRTFs seem to affect the SNR more than the binaural unmasking component.

The underlying reason for the differences in intelligibility across HRTFs is acoustic in nature, and thus driven by differences in the anatomy of the body, head and ears. Nevertheless, we cannot rule out differences that are due to the measurements of the HRTFs. A total of 96 HRTFs from two HRTF databases were used in the current study, and the results across the two databases were similar, reducing the likelihood that the observed effects were due to measurement uncertainties. The databases are likely to cover a diverse range of people, however, it remains unclear if conclusions can be drawn for a worldwide population. The artificial head, which is supposed to reflect an average across the population, for example, showed results slightly above the median predicted intelligibility in the single interferer condition. Thus, the artificial head might not reflect an average head, the databases do not cover an average of the population, or both. However, with symmetric interferers the difference diminishes.

For applications like virtual reality, it is often not possible to use individual HRTFs, thus non-individual HRTFs from artificial heads or databases need to be employed. When choosing a non-individual HRTF for a task that involves speech intelligibility, e.g. a cocktail party situation, it is important to keep in mind which HRTF to choose, as individual characteristics of each HRTF seem to have an impact on the speech intelligibility, as shown in this study. In the case of the CIPIC database, it has been shown that the artificial head HRTF is a good average for most human HRTFs. Thus, using a artificial head HRTF seems to be a viable option when no individual HRTFs are available.

Throughout this study it has been shown that there is an effect of HRTFs on modeled speech intelligibility, however, it remains unclear if listeners are affected by the differences in cue availability or if the auditory system is trained to a specific set of cues. Furthermore, other factors are known to affect speech intelligibility. For example the type of the interferer. Using the current modelling approach assumes static noise interferers with the same spectrum as the target speech. When using speech instead of noise interferers, the speech intelligibility thresholds are commonly increasing (e.g. Carhart *et al.*, 1969) and the SRM is often larger (e.g. Freyman *et al.*, 1999). If speech interferers affect the speech intelligibility with various HRTFs differently than the modelled noise interferers could be investigated in listener studies. Another factor affecting speech intelligibility is room reverberation (e.g. Houtgast *et al.*, 1980; Plomp, 1976). Reverberation has been shown to mask spectral differences in impulse responses (Ahrens *et al.*, 2020; Oreinos and Buchholz, 2015). Furthermore, individual perception differences (e.g. due to cognitive factors or threshold differences) might be larger than the variations across HRTFs and thus mask differences in speech perception as shown in this study. Nevertheless, the modelling approach in this study showed possible differences in cue availability for speech intelligibility with various HRTFs.

## Acknowledgments

We would like to thank Pavel Zahorik and Sebastià Vicenç Amengual Garí for the discussion that led to this manuscript. A.A. was supported by the Centre for Applied Hearing Research (CAHR).

## Notes

### Competing Interest Statement

The authors have declared no competing interest.

